# Neural recordings of continuous speech reveal robust signatures of prediction in second language learners of English

**DOI:** 10.1101/2025.11.10.684289

**Authors:** Craig A. Thorburn, I.M Dushyanthi Karunathilake, London N. Dixon, Ellen Lau, Jonathan Z. Simon

## Abstract

When listening to speech in their native language, speakers use prior context to anticipate upcoming phonemes, words, and concepts, integrating information at the sublexical, lexical, and sentence level. While it has been suggested that late second language learners do not predict to the same extent as native listeners, adequately evaluating this claim requires measurement of predictions at these multiple levels of representation simultaneously in natural speech. We recorded magnetoencephalography (MEG) responses from native Mandarin and Sinhala speakers listening to continuous narrative English speech. We used multivariate temporal response function (mTRF) analysis to investigate whether second language listeners demonstrate the same markers of prediction in neural data as native English speakers listening to the same stimuli. We demonstrate that late second language listeners exhibit strikingly similar responses to native speakers in sensitivity to phoneme surprisal and entropy with respect to sublexical, lexical, and sentence-level context. The few small response differences we observed appear most likely to arise from specific properties of the native languages, rather than general differences between native and second-language listening. These results provide evidence that late second-language listeners indeed leverage prediction in similar ways as native listeners in understanding continuous speech. This suggests that multivariate analyses of neural data from naturalistic listening may be vital in carefully evaluating the differences and similarities in speech prediction across populations.

**Significance Statement:** Much is still unknown about how people listening to a second language predict upcoming words and sounds. Here, we leverage neuroimaging during continuous speech and analyze responses to multiple speech language features in the signal to study the neural encoding of prediction simultaneously at multiple levels of linguistic context. We observe robust encoding of statistical properties tied to prediction at all context levels in second-language learners of English and that responses are strikingly similar between native and second language listeners. Speech language features are encoded similarly in both groups of language learners, with few differences between the native and second language listeners, indicating that second language listeners predict upcoming input similarly to native listeners.

## Introduction

Almost half of the world’s population knows more than one language (Matthews, 2019), with many people acquiring a second language later in life. Yet the neural mechanisms behind second language comprehension are much less studied than those in native language understanding, and many questions remain, particularly how listeners use context to predict upcoming input, a critical aspect of language fluency. When listening to their native language, a listener constantly anticipates what comes next (Ferreira & Chantavarin, 2018), integrating information at a range of levels in the linguistic hierarchy (Brodbeck et al., 2022). However little work has examined how second language listeners integrate these different levels of linguistic context.

Neural and behavioral results typically show differences between prediction in first- and second-language learners^1^ in constrained experimental paradigms, indicating that there may be differences in predictive processes between these two groups. Unlike native speakers, second-language learners may not use certain grammatical cues to restrict their prediction of an upcoming noun, such as the grammatical gender of a determiner (Lew-Williams & Fernald, 2010). Some event-related neural responses to syntactic and semantic violations in second-language listeners may be smaller (Hahne, 2001) or delayed (McLaughlin, Osterhout, & Kim, 2004; McLaughlin et al., 2010) or in fact not present at all (Hahne & Friederici, 2001).

Two possible theories might explain why these differences might be observed. First, it is possible that second language learners have fundamental differences in syntactic parsing and may not represent and use the same information as native speakers during online processing. The Shallow Structure Hypothesis (Clahsen & Felser, 2006a, 2006c, 2006b), suggests that second-language learners rely more on semantic and other non-grammatical cues when listening to speech, as opposed to particular grammatical features.

Alternatively, recent theories suggest that when listening to a second language one may use identical processing mechanisms regardless of language, but the statistical knowledge that has been acquired of a second language is fundamentally different to that of a native language (Kaan, 2014). Differences in statistical knowledge might appear as though listeners are forming predictions using different mechanisms, where in fact they arise due to differing underlying knowledge. Additionally, tasks shown in typical experimental paradigms may be more effortful for second-language listeners, further serving to emphasize the differences between second- and native-language responses.

To address these issues we record neural responses using magnetoencephalography (MEG) while native Sinhala and Mandarin second-language learners of English listen to an audiobook—a more ecological setting than controlled experiments. We leverage multivariate Temporal Response Function (mTRF) analysis (Brodbeck, 2023; Crosse, Di Liberto, Bednar, & Lalor, 2016) to extract responses to prediction of an upcoming phoneme using information at the sublexical, lexical and sentence context level, comparing to the responses of native English listeners. This analysis framework allows us to look at responses across different levels of linguistic context simultaneously across a naturalistic input. Although none of the participants in our study learned English before age 5, they have had substantial exposure to English since then, in principle allowing for them to gain a more ‘native-like’ statistical distribution of the language.

Our results show robust neural encoding of these prediction-related statistical properties using multiple levels of linguistic context in advanced second-language learners. We show that responses are strikingly similar to native listeners at the sublexical and lexical level, with key differences observed at the sentence level. That our results show few differences demonstrates that it is possible for second-language learners to advance to a point where predictive processes essentially mirror those of native listeners. Where we see minor differences is instead between the two second-language learner groups, indicating specific impacts of native language properties on predictive models rather than a general impact of ‘nativeness’. We suggest that the same mechanisms are used during first and second language processing and that task-related effects, knowledge of the statistical environment of the language, and one’s specific native language may play a large role in differences commonly seen in the second language literature.

## Methods and Materials

Second language learners of English listened to an audiobook while MEG responses were recorded during continuous speech listening, using an identical paradigm to one that Brodbeck et al. (2022) used with native English listeners. In order to keep variability across participants to a minimum, while still allowing for comparison between different languages, two specific languages were chosen for this study—Mandarin and Sinhala. These languages were chosen to represent two language families with significant differences from English, from which native speakers were relatively easy to recruit in our local context.

### Data Availability

The raw MEG data from this experiment is available at https://doi.org/10.5061/dryad.wdbrv1636 (Data will be made available upon journal acceptance).

### Participants

Non-native participants were native speakers of Sinhala or Mandarin, had not lived in a primarily English-speaking country for more than 6 months before age 12, and had not learned English before age 5. All language background responses were self-reported. A total of 12 native Mandarin speakers and 12 native Sinhala speakers were included in the reported dataset. Four other Sinhala speakers also participated, but data for three of these participants were excluded when it was later discovered that they did not meet our criteria for late English learners, and the last participant was excluded due to excessive motion during the recording. Native speaker data came from the dataset previously reported by Brodbeck et al. (2022), which included 12 native English speakers.

All subjects self-reported no neurological or hearing impairments and all except 1 native speaker were right-handed according to the Edinburgh handedness inventory (Oldfield, 1971). Each participant gave informed written consent and procedures were approved by the University of Maryland Institutional Review Board. Each subject received compensation for their participation in the 2-2.5 hour experiment.

### Stimuli

Participants listened to approximately 50 minutes of the audiobook ‘The Botany of Desire’ by Michael Pollan (Pollan, 2002). The book was split into 11 segments (205— 331 seconds) identical to those in Brodbeck et al. (2022). To ensure that participants processed the excerpts for meaning, they were presented in chronological order.

### Procedure

Before scanning commenced, participants were presented with an English exposure questionnaire. The questionnaire was a short survey that asked about participants’ exposure to the English language, length of residency in the United States, and comfort with English. Questions on this survey and responses are reported in Table 1.

**Table 1.** Profiency Profile [Mean (Standard Deviation)] in years for all 24 analyzed participants.

| Age | Sinhala | Mandarin | All |
| --- | --- | --- | --- |
| **Language Exposure Questions** |  |  |  |
| At what age did you first learn English? | 6.0 (2.1) | 7.5 (3.3) | 6.7 (3.2) |
| At what age did you feel proficient in English? | 18.5 (5.0) | 18.4 (3.5) | 18.4 (4.4) |
| How long have you lived in the US? | 2.4 (1.2) | 3.7 (2.7) | 3.0 (2.1) |
| At what age did you spend more than 3 hours a day in an English speaking environment? | 14.4 (5.6) | 15.8 (3.9) | 15.1 (4.9) |

Participants then lay in the scanner and first took part in two brief scans consisting of a 60 second recording of resting state activity and a tone counting task where participants heard 100 jittered tones. Participants were asked to either keep their eyes open or closed for the entire duration of these tasks and during audiobook listening, and avoid excessive blinking or movement.

Participants then listened to the audiobook in 11 segments with short breaks in between. During each break, participants were asked 3-4 comprehension questions to encourage them to actively attend the content of the story. No participants were excluded on the basis of their answers to these questions.

### Data Acquisition and Preprocessing

Neural activity was recorded using a whole-head MEG system (KIT, Kanazawa, Japan) with 157 axial gradiometers inside a magnetically shielded room (Vacuumschmelze GmbH & Co. KG, Hanau, Germany) at the University of Maryland, College Park. Sensors (15.5 mm diameter) are uniformly distributed inside a liquid-He dewar, spaced ∼25 mm apart, and configured as first-order axial gradiometers with 50 mm separation and sensitivity better than 5 fT Hz*^−^*^1^*^/^*^2^ in the white noise region (>1 kHz). Data were recorded with a sampling rate of 1 kHz and online filtered with a 200 Hz low-pass filter and a 60 Hz notch filter. Auditory stimuli were delivered through foam pad earphones inserted into the ear canal at a comfortably loud listening level.

Data were processed and analyzed using MNEPython (Gramfort et al., 2014) and Eelbrain (Brodbeck et al., 2021) with additional TRF-Tools helper scripts (Brodbeck, 2023; Brodbeck et al., 2022), closely mirroring the analysis of Brodbeck et al. (2022). Flat channels were automatically detected and excluded, spatiotemporal signal space separation (Taulu & Simola, 2006) was applied to remove artifacts, and data was filtered between 1-40 Hz with a zero-phase FIR filter (mne-python 0.20 default settings). Independent component analysis (ICA) (Bell & Sejnowski, 1995) was then used to remove additional artifacts of eye blinks and heart rate and from the MEG signal based on visual inspection. A further 1-10 Hz bandpass filter was applied.

Prior to recording, participants’ head shape was digitized (Polhemus 3SPACE FASTRAK) and head position in the scanner was measured using 5 marker coils— 3 placed on the forehead and two placed 1 cm anterior from the tragus on each ear. Head position was measured at the beginning and end of the experiment, and the two measurements were averaged for coregistration purposes. The head shape of each participant was fit to the FreeSurfer *‘fsaverage’* average brain (Fischl, 2012) using an average scaling technique and this coregistration was used to create an inverse function such that activity could be mapped to virtual source dipoles across a common cortical surface. A source space was generated using fourfold icosahedral subdivision of the white matter surface, with source dipoles oriented perpendicularly to the cortical surface. Regularized minimum l2 norm current estimates (Dale and Sereno, 1993; Hämäläinen and Ilmoniemi, 1994) were computed for all data using an empty room noise covariance (*λ* = 1/6).

Cortical regions were determined using the ‘*aparc*’ freesurfer parcellation (Desikan et al., 2006). Preliminary analysis took place over the entire cortical surface, excluding the occipital lobe, insula and midline structures. Further analysis was restricted to an a priori ROI that includes bilateral superior temporal areas— specifically the *superiortemporal* and *transversetemporal* labeled ‘*aparc*’ parcellations. This a priori ROI was chosen as it includes areas that primarily deal with speech processing, and has shown robust previous encoding in native language listeners for the predictors used in the current analysis (Brodbeck et al., 2022).

### Experimental Design and Statistical Analysis

The continuous neural data was analyzed using an mTRF analysis technique (Brodbeck et al., 2022; Brodbeck, 2023) where the signal is deconvolved with various speech language features to uncover neural encoding of prediction throughout the audiobook.

Under this method, it is assumed that some of the neural signal can be generated by convolving a particular response function with the value of each speech language feature at each point in time. These features can either be single impulses—such as at the onset of a phoneme—or continuous features such as the acoustic envelope of the stimulus.

The deconvolution allows the generation of specific temporal response functions (TRFs) that model the neural response to the presence of each feature. The deconvolution is performed individually for each virtual source dipole, allowing a spatial measure of how significantly each predictor is able to explain the neural signal across the cortical surface.

The models and predictors used follow the methodology from Brodbeck et al. (2022).

**Phonetic Transcription** A phonetic transcription of the audiobook was obtained using a combination of the CMU pronunciation dictionary and the Montreal forced aligner (McAuliffe, Socolof, Mihuc, Wagner, & Sonderegger, 2017), which was also used to align the phonetic transcription with the acoustic stimulus. Additional manual adjustment was performed to align acoustic data with the transcriptions.

**Acoustic Predictors** To absorb any activity in the neural signal that can be better described by low-level acoustic properties of the stimulus, predictors based on gammatone envelopes and gammatone onsets are included in all models, with 8 log spaced frequency bands between 10 and 5000 Hz (Brodbeck et al., 2022). In addition, to account for neural signals that correspond to the segmentation of phonemes and for any variance that can be represented purely with phoneme boundaries, two predictors are included, corresponding to phoneme and word onset. The word onset predictor is comprised of an impulse for every word onset and phoneme onset is comprised of an impulse at each phoneme onset that is not also a word onset (to avoid colinearity issues). Therefore, any neural responses described by each of the speech language features below can be attributed to the specific predictor that it encodes and not simply because there is a word or phoneme onset.

**Context Models** Models of surprisal and entropy of phonemes at three different context levels are used to study the integration of context information at multiple levels of the linguistic hierarchy, as described in previous work.

### Sublexical Context Model

Sublexical predictors are calculated as the probability of encountering a particular phoneme, given the previous 4 phonemes heard, yielding a 5-gram phoneme prediction model. These values are generated by training a 5-gram model on the SUBTLEX-US corpus (Brysbaert & New, 2009). Two values of this distribution are used as features in the TRF model: phoneme surprisal, and phoneme entropy. Surprisal is calculated as in (1), where *phoneme_k_* is the current phoneme and *context* include the previous 4 phonemes

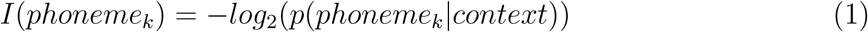

Phoneme entropy is defined as the average expected surprisal of the following phoneme (2). Here the context is the previous 3 phonemes and current phoneme *phoneme_k−_*_3;_*_…k_*, and *E* represents the entire inventory of phonemes in the language.

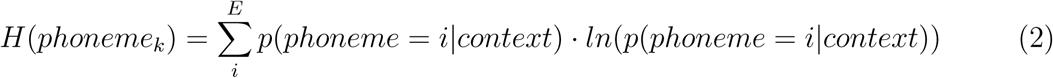

### Lexical Context Model

For the lexical context model, the same equations apply, but starting from the onset of a word. The probability of a given word is the initial probability of the word calculated from word frequency extracted from the Corpus of Contemporary American English (Davies, 2017). At each subsequent phoneme, the probability of any word that is not consistent with the phonemes already heard is set to zero, and the distribution is normalized. The probability of a phoneme is calculated given the probability of every possible word completion.

In addition to the measures described above, an additional statistical measure can be calculated as the entropy over the distribution of words. This is calculated in the same way as phoneme entropy above, except with word probability instead of phoneme probability.

Here *phoneme_j,k_*represents the *k*-th phoneme in the *j*-th word and *W* represents all words in the corpus.

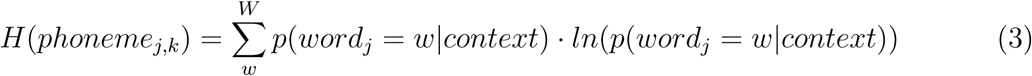

### Sentence Context Model

For the sentence context model a 5-gram model is trained on the Corpus of Contemporary American English (COCA) (Davies, 2017) with KenLM (Heafield, 2011), where word probability is calculated given the previous 4 words seen. For sentence context models, the same measures above are calculated, however instead of the probability of a word proportionate to its frequency, it is equal to its probability given the 5-gram model.

**Models** As in Brodbeck et al. (2022), deconvolution and statistical analysis were performed with Eelbrain (Brodbeck et al., 2021) and additional scripts available at https://github.com/christianbrodbeck/TRF-Tools (Brodbeck, 2021).

mTRFs were computed independently for each subject and each virtual current source (Brodbeck et al., 2018b; Lalor et al., 2009). mTRFs were generated from a basis of 50 ms wide Hamming windows centered at *T* = [−100*, . . .,* 1000) ms. For estimating mTRFs, all responses and predictors were standardized by centering and dividing by the mean absolute value.

For estimation using fourfold cross-validation, each subject’s data were concatenated along the time axis and split into four contiguous segments of equal length. The mTRFs for predicting the responses in each segment were trained using coordinate descent (David et al., 2007) to minimize the l1 error in the three remaining segments. For each test segment there were three training runs, with each of the remaining segments serving as the validation segment once. In each training run, the mTRF was iteratively modified based on the maximum error reduction in two training segments (the steepest coordinate descent) and validated based on the error in the validation segment. Whenever a training step caused an increase of error in the validation segment, the TRF for the predictor responsible for the increase was frozen. Training continued until the whole mTRF was frozen. The three mTRFs from the three training runs were then averaged to predict responses in the left-out testing segment.

For each virtual source dipole in each subject, mTRFS were generated independently. The full mTRF model includes acoustic and phoneme onset features as well as surprisal, cohort entropy and phoneme entropy calculated using the lexical and sentence context levels and phoneme entropy and surprisal calculated using the sublexical context level (cohort entropy is not defined at the sublexical context level). To calculate the predictive power of a feature or context level, a reduced model is run including all features in the full model, except the feature(s) under investigation. We construct a ‘model quality index’ defined as 1 − (*l*1(residuals)*/l*1(*y*)), where *y* is the continuous neural signal, giving an indication of how good a model is fit to the data. The increase in model quality index from the reduced model to the full model yields a conservative estimate of the improvement that this specific feature or set of features contribute to the model fit.

The improvement for each level of linguistic context is measured by calculating this difference in model quality index for each individual virtual source dipole and participant.

We smooth this difference using a Gaussian window and apply a mass-univariate one-sample t-test with Threshold Free Cluster Enhancement (TFCE) (Smith & Nichols, 2009).

To test differences of lateralization, we morph the cortical map of differences in predictive power to the FreeSurfer *‘fsaverage’* symmetrical average brain (Greve et al., 2013) and map from the right hemisphere onto the left hemisphere. We then once again use a mass-univariate paired t-test with TFCE to determine whether there is a statistical difference between responses in the two hemispheres for each language group. Additionally, we calculated a lateralization index to measure the lateralization of a specific speech language feature using the formula: *LI* = *R/*(*L* + *R*), where *R* and *L* are the predictor strengths in each hemisphere.

To measure differences in the encoding of each predictor and to investigate differences between each language group, we conduct tests using the improvement in model quality index for each feature over all virtual source dipoles in our STG ROI of Superior Temporal Gyrus. For these tests we take the average predictive power over all virtual source dipoles within this ROI for each predictor and participant. We then conduct a one-sample t-test over these data to determine whether the model improvement is significantly above zero.

We additionally run a linear regression to test whether the mean improvement across participants varies significantly across the 3 languages. A regression is run for each speech language feature and hemisphere with language as a fixed effect. For any models where language is a significant factor, we perform further t-tests between each language pair: English-Mandarin, Sinhala-Mandarin, Mandarin-Sinhala.

Finally, we calculate study differences in the TRF for each predictor itself using cluster-based permutation testing. Clusters of differences between native and second language learners in both space (virtual source dipoles in STG) and time are tested for significance using permutation analysis. We additionally perform a cluster-picking analysis, where we extract the size and time point of each peak in spatially averaged TRF for each participant for every predictor across the two language groups. Performing independent t-tests yields no significant differences between the size and location of each peak across all speech language features. We therefore do not report these analyses in this paper.

## Results

### All language listeners show strong cortical prediction-related responses

We first show that there is significant encoding of prediction at each level of the linguistic hierarchy in both groups of second language listeners. This replicates the results of Brodbeck et al. (2022) which similarly demonstrate significant encoding in native speakers. Each of the levels of context we use — sublexical, lexical, and sentence — uniquely contributed to explaining the neural data, in each of the three speaker groups, with significant improvement in model quality over a model that omits features at that specific context level (Table 2). Figure 1 shows the cortical maps of this improvement for each language group and context level. As expected, much of the explained variance lies in STG for every level of context, across groups. Although it is of interest that these effects appeared to additionally extend more inferiorly to MTG for sentence-level context, we restrict further analyses in the current paper to our a priori STG ROI as this facilitates the comparisons across context levels and groups.

**Figure 1.**
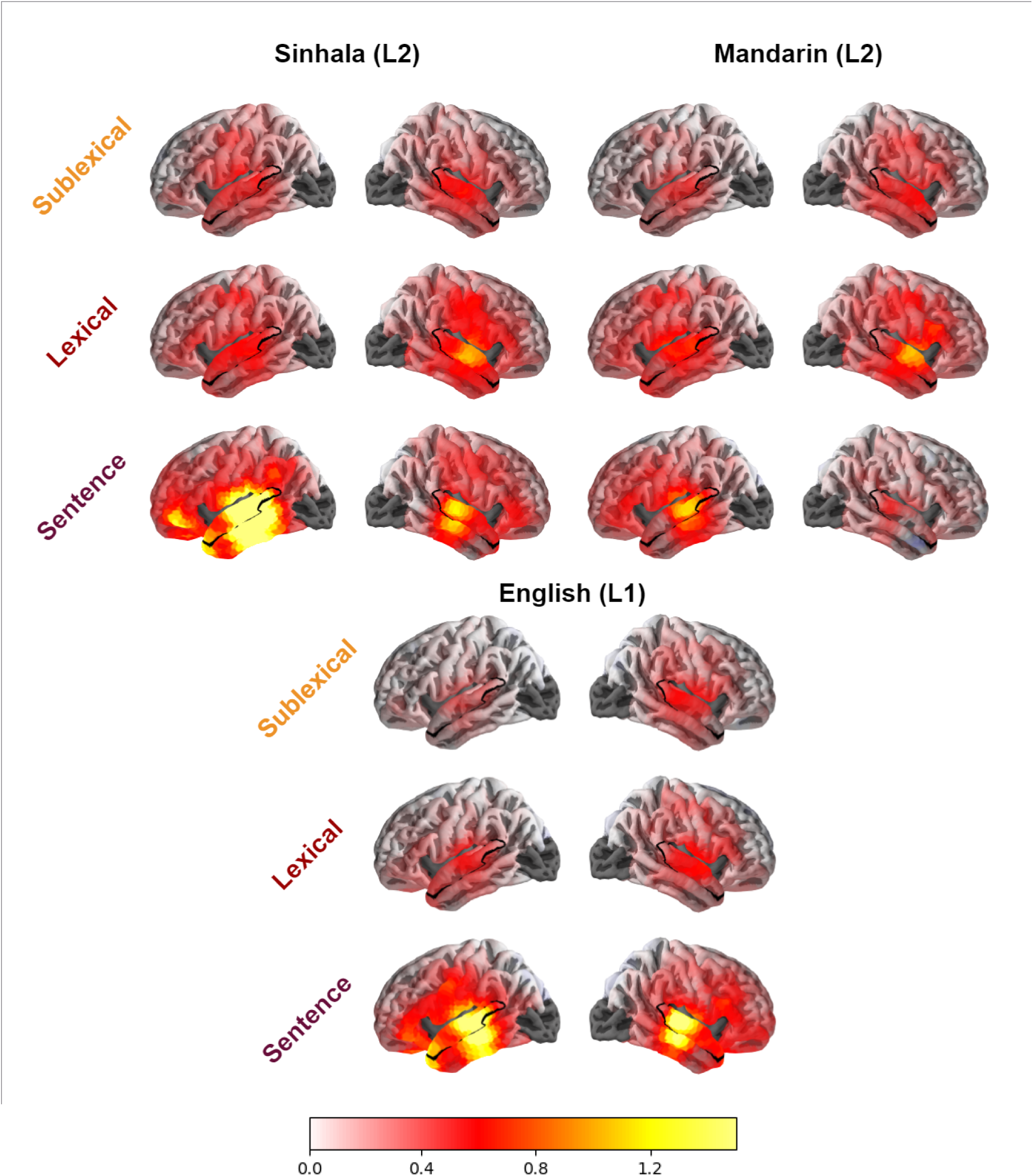
Model improvement for each context level mapped over cortex in native speakers of each of the three languages. Significance values reported as mass univariate t-Tests over cortex are given in Table 2. The inclusion of each of the context levels improves the model quality index across all three languages.

**Table 2.** Mass Univariate Statistics for model improvement at each context level for all cortex for L1 and L2 listeners. Statistical values shown are for the most significant cluster across cortex in each test, with tmax as the t value for the largest difference cluster observed across space and time, with the p value calculated through cluster-based permutation testing.

| Mass Univariate Tests Over Cortex |  |  |  |  |  |  |
| --- | --- | --- | --- | --- | --- | --- |
|  | L1 |  |  | L2 |  |  |
|  | tmax(11) | p |  | tmax(23) | p |  |
| Sentence | 12.2 | <0.001 | *** | 6.2 | <0.001 | *** |
| Lexical | 11.2 | <0.001 | *** | 9.4 | <0.001 | *** |
| Sublexical | 7.8 | 0.002 | ** | 7.7 | <0.001 | *** |

**Table 3.**
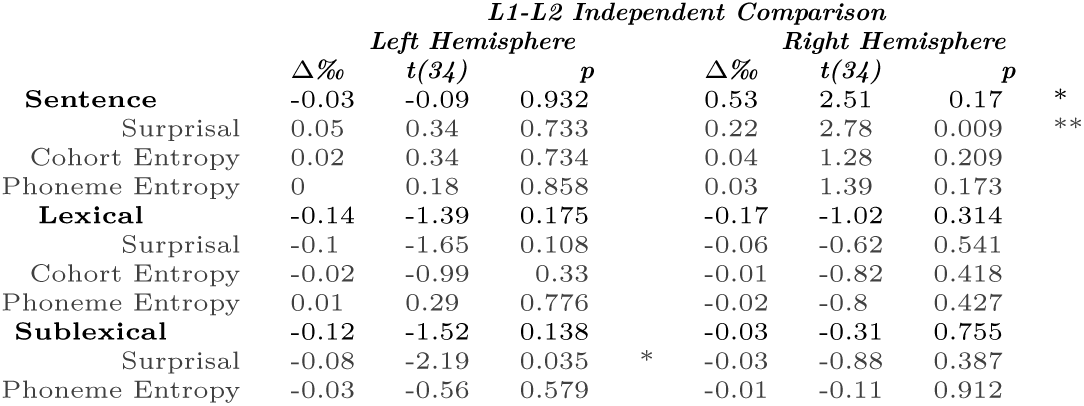
Simple t-tests comparing average model prediction improvement between L1 and L2 groups. Improvement for each predictor calculated for each participant averaged over STG ROI and then independent two-sample t-test ran between different language groups. We see only sublexical surprisal, sentence surprisal and sentence predictors overall show significant differences. For the most part, there are no significant differences between L1 and L2 speakers.

Within this ROI we further break down these effects by individual predictor — surprisal, cohort entropy and phoneme entropy (Figure 2), showing the improvement in model quality individually for each listener, compared to a model that does not include this feature. Each of these improvements differs significantly from zero in a simple one-sample t-test (Table 4), showing that not only does each context level explain significant variance when all individual features at the level are taken together, but that each individual feature uniquely explains some variance in responses within STG.

**Figure 2.**
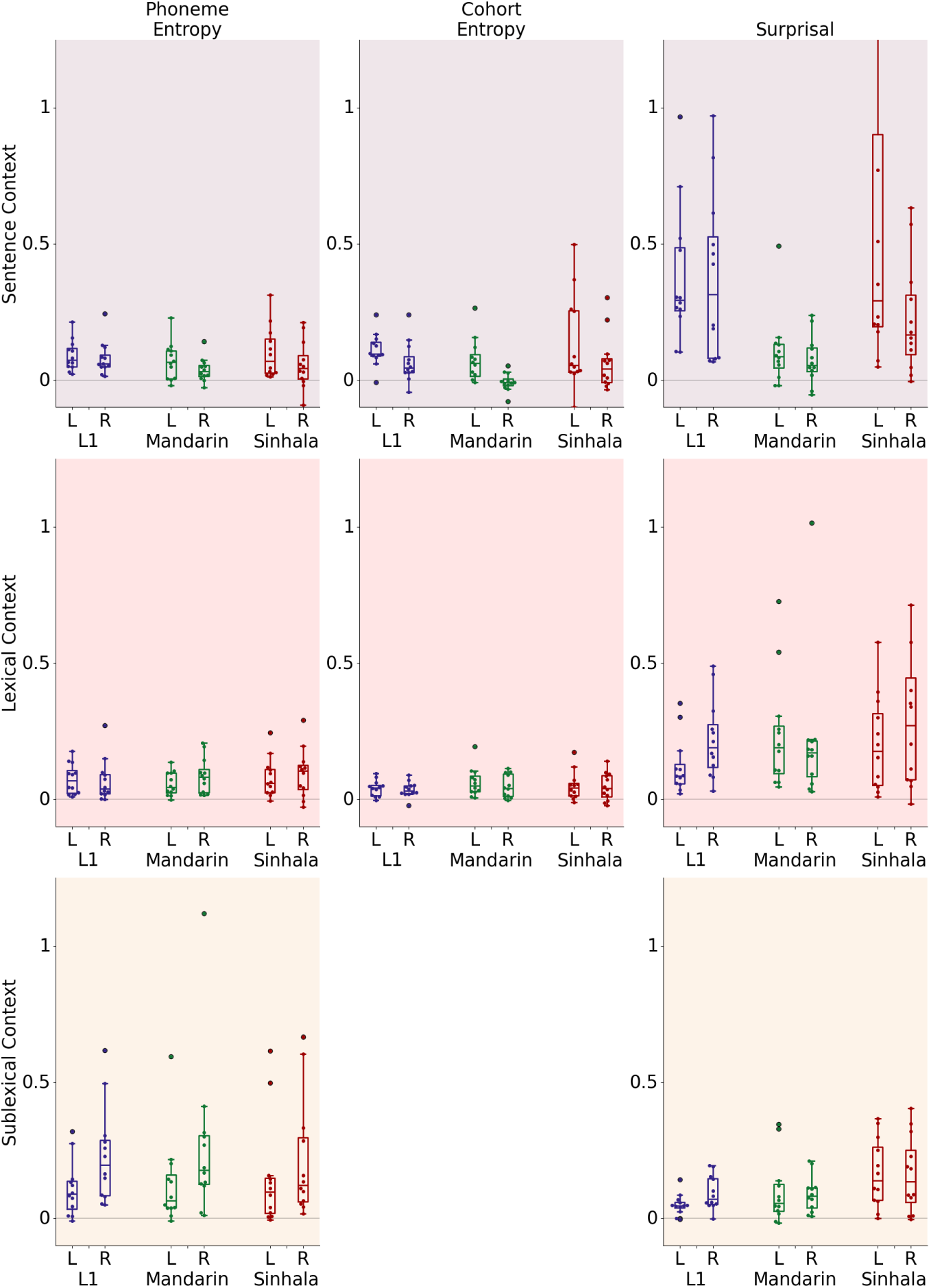
Model Improvement for each predictor averaged across source space. Each datapoint represents an individual participant with analysis restricted to STG. Results are expressed as a percentage of the highest explained variability of the full model across all dipoles in both language groups. *All improvements are significantly above zero on a paired t-test on the average model improvement across the STG with specific tests reported in Table 4.* Cohort entropy is not defined for sublexical information.

**Table 4.**
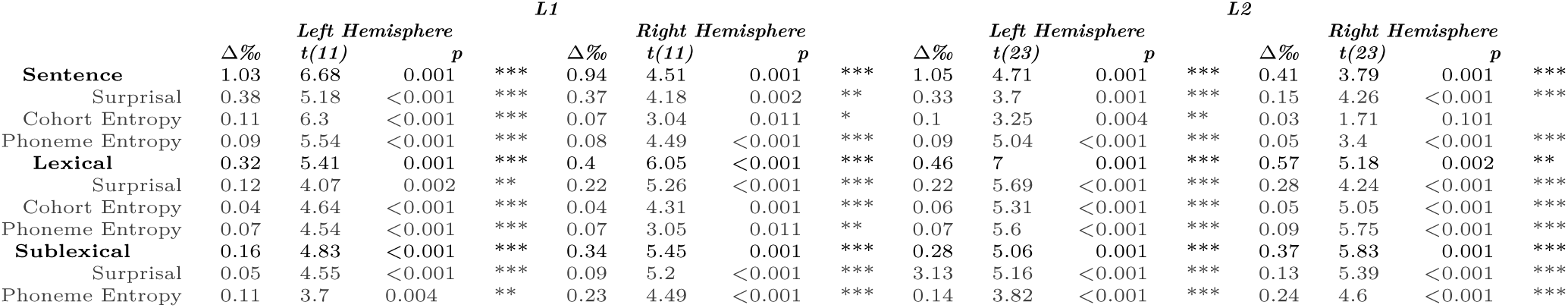
Simple t-tests for model prediction improvement for each participant, averaged over STG ROI (ie. each participant has single data point for each model which is averaged over all sources across the STG). Each predictor in each model significantly improves the fit of the model in at least one hemisphere (only one predictor—right-hemisphere sentence cohort entropy did not reach significance in L2 speakers).

**Table 5.**
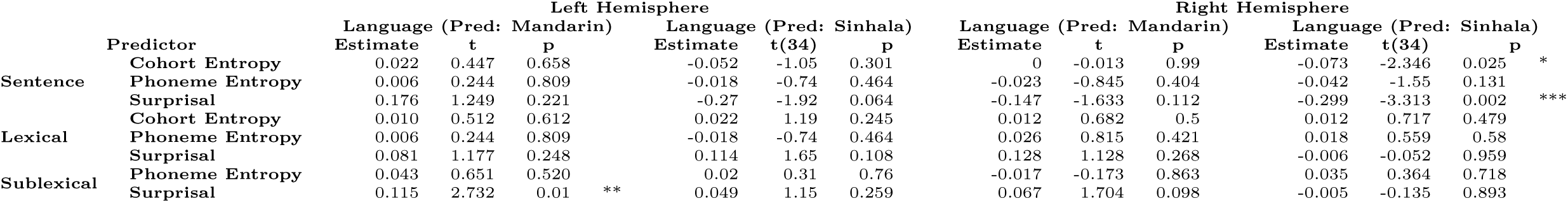
Output of linear regression to test contribution of language to the model improvement of each predictor within STG. Estimates, t-tests and p-values shown for individual regressions for each predictor and hemisphere, with language as a predictor in the model. Baseline value is encoded as English. For models which show a significant contribution of language to the model improvement, further individual t-test comparisons between each language pair are shown in Table 6.

### Responses between first and second language learners are closely aligned, with some minor differences

While each predictor significantly captures the neural signal across all native languages, one of our primary goals is to investigate whether these data differ significantly between the two second language groups and whether they differ from native speakers. To achieve this we run linear regression models for each feature, with listener native language as a predictor in the regression (Table 4). For any of these models which yield language as a significant predictor, we run three t-tests to compare each language pairing (Table 6).

**Table 6.** T-Test comparisons between each language pairing, for any predictor and hemisphere where one language is a significant predictor.

|  | Sinhala-Mandarin |  |  | Mandarin-English |  |  | Sinhala-English |  |  |
| --- | --- | --- | --- | --- | --- | --- | --- | --- | --- |
|  | t(23) | p |  | t | p |  | t(23) | p |  |
| Sentence Cohort Entropy RH | 2.316 | 0.030 | * | 3.074 | 0.006 | *** | 0.011 | 0.992 |  |
| Sentence Surprisal RH | 2.338 | 0.029 | * | 3.208 | 0.004 | *** | 1.372 | 0.184 |  |
| Sublexical Surprisal LH | 3.167 | 0.002 | *** | -1.327 | 0.198 |  | -3.019 | 0.006 | *** |

For most speech features, we found that language group was not a significant predictor of model improvement. For three features however, the linear regression model did find a significant effect of language: sentence cohort (right hemisphere only), sentence surprisal (right hemisphere only), and sublexical surprisal (left hemisphere only).

Follow-up independent samples t-tests indicated that the right-hemisphere differences reflected smaller right-hemisphere sentence context effects for native Mandarin speakers compared to the other two language groups. The left-hemisphere sublexical surprisal difference appeared to be driven by a more robust left-hemisphere sublexical surprisal effect in Sinhala compared to the other two language groups. Similarly for sublexical surprisal, we observe significant differences in the Sinhala-Mandarin and Sinhala-English pairings, suggesting that Sinhala differs from the other languages for this predictor.

In sum, detailed comparisons of the individual context feature responses in each language group revealed few differences overall. The three cases in which differences were observed appear to be attributable to individual language properties rather than a systematic difference between native and second language listeners.

### Hemispheric Lateralization of Effects

We next investigated potential differences in lateralization by calculating a lateralization index for each feature and context level, shown in Figure 3. In Brodbeck et al.’s 2022 native speaker data, there was a tendency towards slightly stronger right hemisphere effects of lexical and sublexical context. Here, mass univariate t-tests of lateralization (calculated as in Brodbeck et al., where cortical responses are mapped to a common source space and compared between hemispheres), indeed indicate a statistically significant lateralization effect for sublexical surprisal (Table 6). Both groups of non-native speakers showed a similar rightward trend numerically for sublexical phoneme entropy, although these lateralization effects were not significant. For lexical surprisal, Sinhala speakers showed a significant lateralization effect, but Mandarin speakers did not. We ran similar linear regressions as above, over the lateralization index calculated for each feature and hemisphere. No differences between language groups are revealed in these tests which average across spatial information in each hemisphere, suggesting that there are no robust or large differences between each of our languages, nor between native and second language listeners.

**Figure 3.**
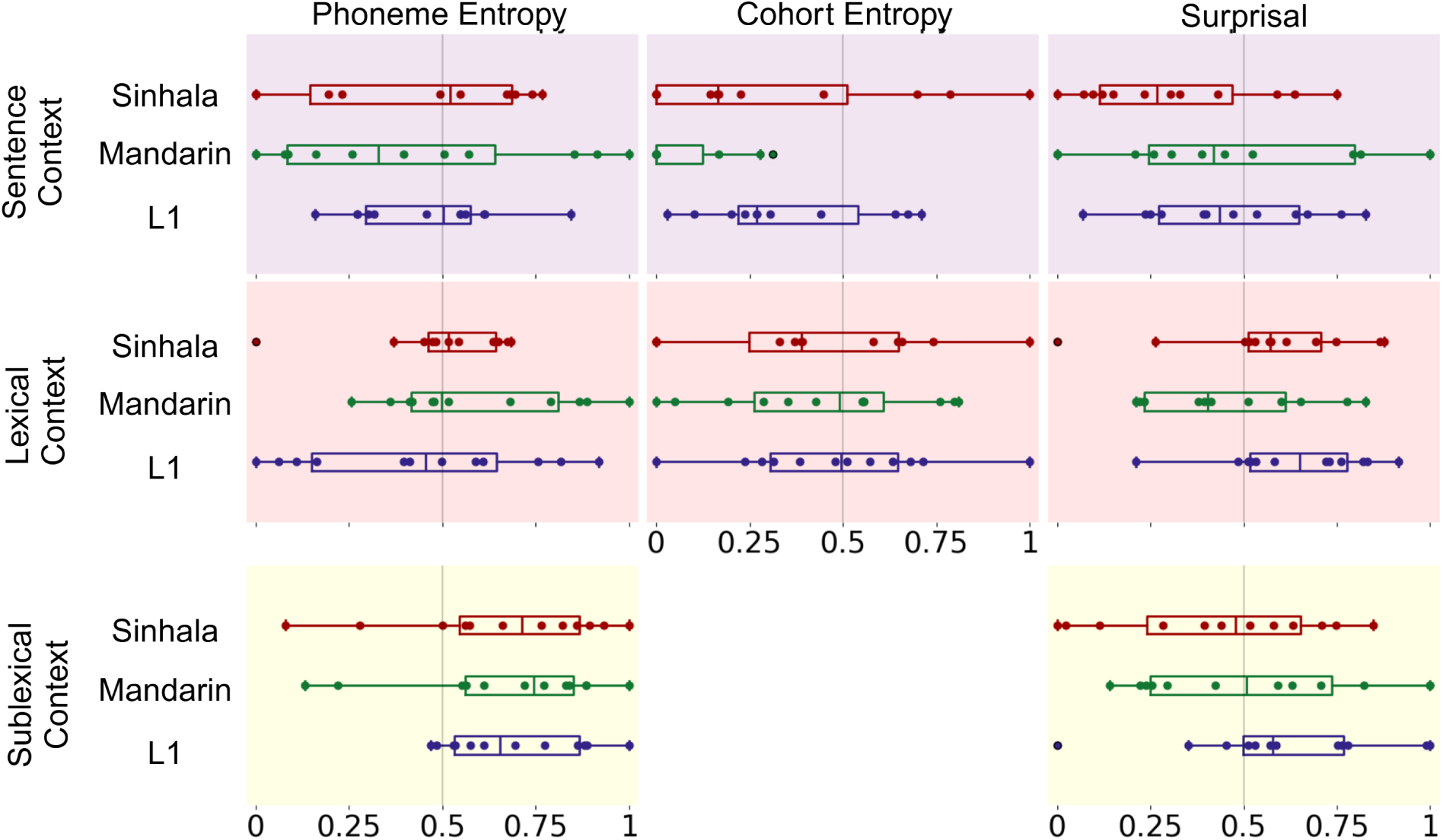
Lateralization Index calculated over model improvement index shown in Figure 2. A value of 0 indicates that all improvement from speech language feature inclusion arises in left hemisphere, 1 indicates right hemisphere. A value of 0.5 indicates equal improvement in right and left hemisphere. Mass univariate tests of lateralization are shown in Table 7 when sources are mapped to common source space. Those tests are reported on this figure and not significant unless reported. Lateralization index across sentence context improvements. Only in English native speakers is significant lateralization for any predictor observed.

**Table 7.**
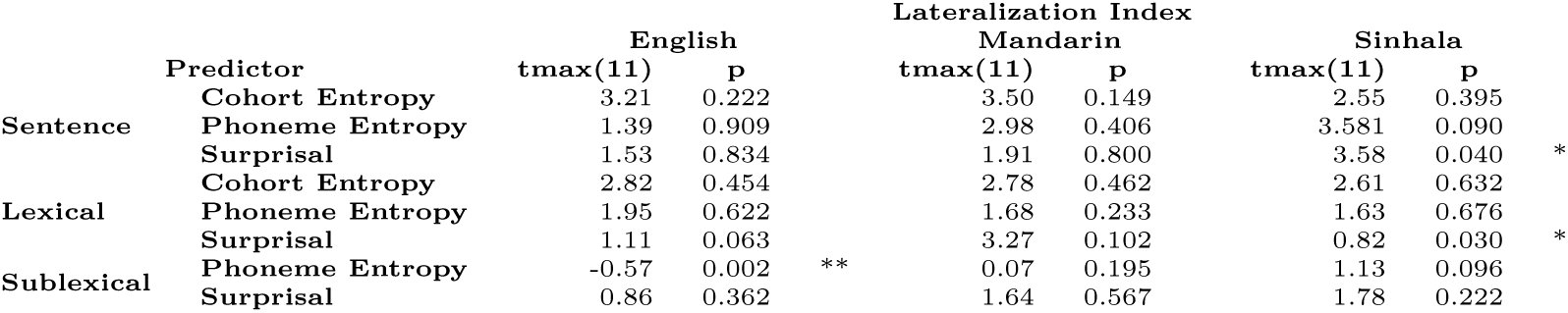
Mass univariate t-Tests comparing model improvement in each hemisphere for L1 and L2 speakers. Responses are mapped to common source space before t-test is run.

**Table 8.** Linear Regression models for the lateralization index of each predictor, where language is used as a predictor. English provides baseline encoding of intercept. For none of our speech language features is language a signfiicant predictor of the lateralization index within this analysis.

|  | Predictor | Lateralization Index |  |  |  |  |  |
| --- | --- | --- | --- | --- | --- | --- | --- |
|  |  | Language (Pred: Mandarin) |  |  | Language (Pred: Sinhala) |  |  |
|  |  | Estimate | t(34) | p | Estimate | t(34) | p |
| Sentence | Cohort Entropy | -0.341 | -1.507 | 0.141 | -0.29 | -1.28 | 0.209 |
|  | Phoneme Entropy | 0.156 | 1.056 | 0.299 | 0.238 | 1.616 | 0.116 |
|  | Surprisal | -0.067 | -0.353 | 0.726 | -0.161 | -0.85 | 0.403 |
| Lexical | Cohort Entropy | -0.269 | -1.217 | 0.232 | -0.298 | -1.35 | 0.187 |
|  | Phoneme Entropy | 0.156 | 1.056 | 0.299 | 0.238 | 1.62 | 0.116 |
|  | Surprisal | -0.180 | -1.036 | 0.308 | -0.21 | -1.21 | 0.236 |
| Sublexical | Phoneme Entropy | -0.021 | -0.201 | 0.842 | -0.026 | -0.25 | 0.802 |
|  | Surprisal | -0.014 | -0.101 | 0.921 | -0.082 | -0.58 | 0.567 |

### Temporal response functions for each predictor are closely matched across language groups

Given the consistency of responses in the two groups of second language learners so far, the temporal response functions themselves are aggregated to analyze timing differences in the temporal response functions between L1 and L2 listeners. As shown in Figure 4, temporal response functions are also closely matched across L1 and L2 speakers, demonstrating that not only is the amount of neural activity that correlates with each predictor consistent across first and second language learners, but even the timecourse of the response is also closely matched between the two groups.

**Figure 4.**
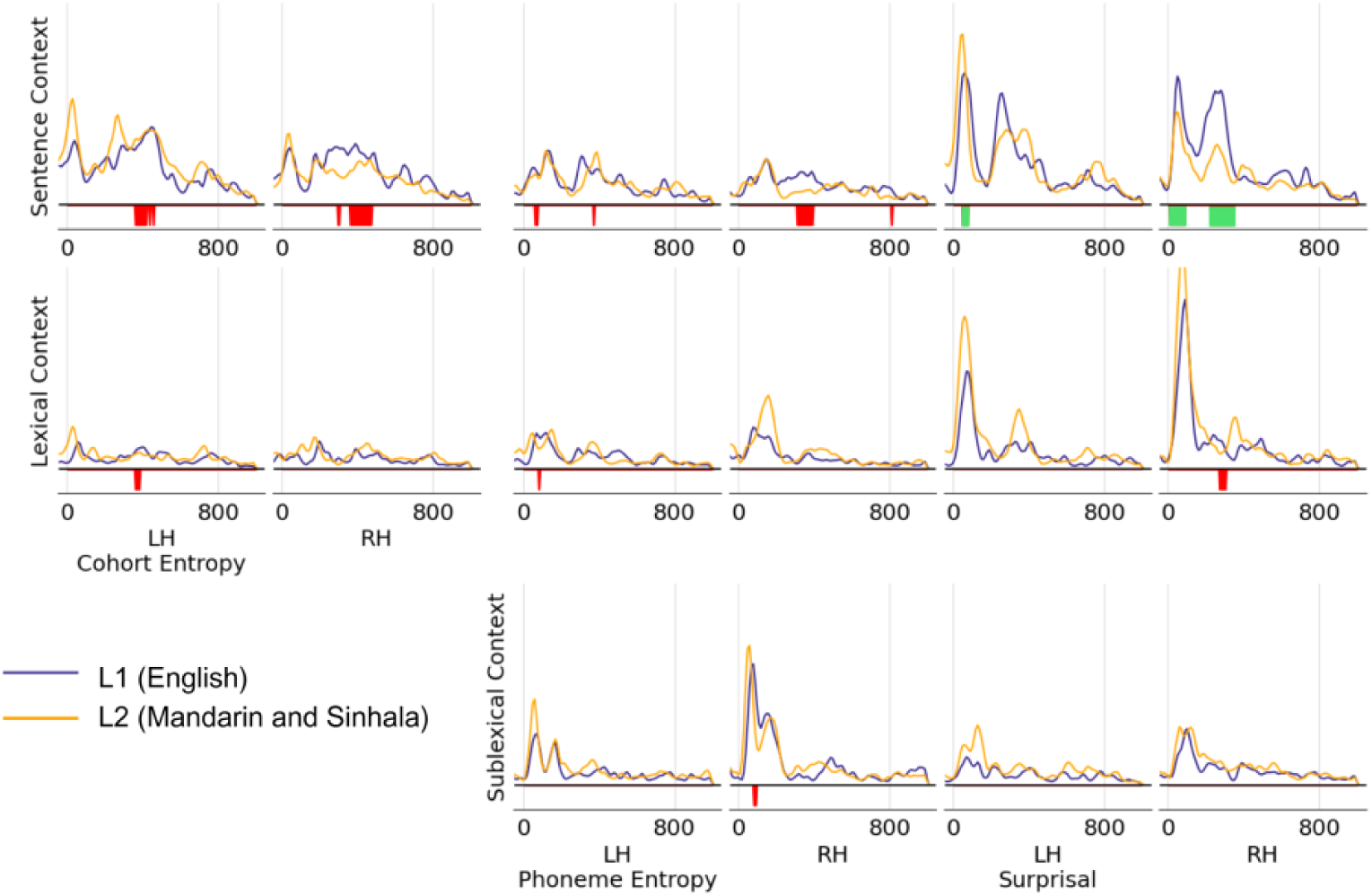
Temporal Response restricted to virtual source dipoles in STG. Between the two listener groups, all dipoles in STG showed significant model prediction improvement for at least one predictor, so these response functions are not masked by specific dipoles within STG. A cluster-based permutation test was ran over all virtual source dipoles in STG—significance bands represent timepoints where at least 2% (red) or 5% (green) of dipoles in the ROI were members of a cluster where the two language groups differ from each other significantly. Few differences between native and second-language listeners are observed across all speech language features, with sentence level differences consistent with the differences found in model improvement analysis.

A clustering analysis reveals some small differences in the response function between native and second language listeners. This is shown by the significant bands illustrated in Figure 4. While the cluster analysis reveals some differences, it is important to note that these clusters only appear spatially in a small number of virtual source dipoles. Only three clusters have a spatial extent that includes 5% or more of the total virtual source dipoles within the corresponding ROI—all of these appearing for sentence surprisal predictor. The largest difference clusters are found in sentence cohort entropy and surprisal in the right hemisphere, overlapping with the differences between languages observed in prior analysis.

Additionally, peak analysis focusing on the location and size of each peak in the TRFs reveals no significant differences between the two language groups in either of these measures for any of the predictors tested.

## Discussion

In this study, we recorded MEG responses from second-language learners of English during naturalistic continuous listening to an audiobook in English and, using mTRF analysis, uncovered neural responses to prediction at multiple levels of the linguistic hierarchy. Our results show robust encoding of prediction using sentence, lexical, and sublexical context in the continuous MEG neural signal. We see that each speech language feature of phoneme entropy, cohort entropy and surprisal, at all context levels, significantly increases the mTRF model fit to the neural data in second language learners, with responses localized to superior temporal regions that are associated with speech processing. Not only do second language learners predict upcoming input at various levels of linguistic context, but these responses are very similar to those of native speakers. MEG recordings from native listeners encode similar information, and there are few statistically significant differences between the two groups. While the participants in this study have reached a high level of proficiency in English, none had learned English before age 5, and were not exposed to more than 3 hours of English a day until age 14. Therefore despite having advanced proficiency at the time of scanning, they clearly meet any standard definition of second-language learners.

These results show strong evidence that prediction in second language learners closely aligns with native speakers, contrasting with prior neural studies showing differences in event-related responses to speech errors between native and second language listeners (McLaughlin et al., 2010, 2004; Hahne & Friederici, 2001; Hahne, 2001). One explanation for these prior results has been the Shallow Structure Hypothesis proposing that second language learners have a reduced ability to use grammatical information during prediction. However, if this were the case, one would not expect second language learners to show robust responses at different linguistic context levels, especially the sentence-level, in contrast to what we observed in the current study. Our results indicate that the neural responses from second language learners encode prediction using sentence-level context information, which likely includes grammatical information tested in prior studies.

Our data aligns more closely with the theory posited by Kaan (2014) who argues that second language learners use the same processing mechanisms as native speakers but form alternative predictions due to differing underlying statistical knowledge. This statistical knowledge could include phoneme transition probabilities and word frequency and transition probabilities—precisely aligned with the metrics tested in the current study. Although none of the participants in our study learned English before age 5, they have had substantial exposure to English since then, in principle meaning they possess a more ‘native-like’ statistical distribution of English. In previous ERP studies, differences may be more prominent as the specific contexts tested may be those where second language learners have not gained sufficient statistical knowledge that mirrors that of native speakers. Kaan (2014) has also argued that researchers may not have been able to see similar predictions between advanced learners and native speakers due to task-based differences. Controlled prediction studies may involve tasks that are more challenging for native speakers, and the inclusion of sentences that violate syntactic and semantic constraints may influence the use of prediction more generally through the study. By testing in a continuous speech paradigm with only syntactically and semantically well-formed input, we ensure that the effects of task difficulty are minimized, providing a more ecologically valid view on what differences might exist between the two language groups.

While our results show little differences between the two language groups across most of our analyses, there are a few measures where we do observe important differences — namely encoding of sentence level predictions and lateralization for a handful of predictors. For sentence context-level effects, our tests reveal that Mandarin listeners in particular appear to show significantly different encoding of this predictor in the response compared to English and Sinhala listeners. Similarly, Sinhala speakers appear to show significantly different encoding to English and Mandarin speakers in the amount of encoding of sublexical surprisal. Although these results are not conclusive, they suggest that individual differences may arise from specific native language transfer effects more than a general difference between native and non-native language processing.

While not statistically robust, we observed some numerical trends towards differences in lateralization between participant groups. For sublexical phoneme entropy, we observed significant right lateralization of responses in native-speakers, but not in the second language learners (Table 6), although the differences between the two groups do not reach significance in our linear regression tests (Table 7). Similarly, right lateralization is observed in the Sinhala speakers for both Sentence and Lexical Surprisal, but once again shows no differences between languages. Exploring these potential differences in lateralization will be an important direction for future work with larger sample sizes and may provide insightful results as to the processes recruited in online phoneme prediction.

The discrimination of phonemes that do not exist in one’s native language is known to be challenging for second language learners to acquire, likely leading to differing representations of phonetic information in these listeners. This may lead to the recruitment of additional neural regions, which may lead to the bilateral activity observed.

Analyzing neural responses using continuous speech recordings allows a hypothesis-free way of observing responses that might not be apparent in more controlled experimental environments. While prediction in second language has been a robust field of research, until now most of the data has been from the processing of controlled experimental stimuli. Our dataset, studying responses from longer story segments presented in a continuous manner, brings a unique perspective that may not be observable by other means. We leverage our data to gather results simultaneously across context levels in the linguistic hierarchy, showing prediction using several layers of contextual information simultaneously within one experiment. This allows us to show that not only do second-language learners’ responses reflect native knowledge in general, but specifically that the responses are consistent across prediction using prior sublexical information, lexical information and sentence information. This also lets us see the specific situations where differences exist between second language and native speakers.

Finally, the current work provides a valuable corpus of second language neural data with unique properties, which can be used easily for further study of speech processing in second language learners without the need for additional experiments. While at least one EEG dataset of second language naturalistic listening exists (Di Liberto et al., 2021), in addition to several continuous fMRI datasets (Brennan & Hale, 2019; Bhattasali et al., 2019), this is one of the first datasets of MEG naturalistic listening data for second language learners. MEG allows not only good spatial resolution across the cortical surface, but it also allows for fine temporal resolution to see rapid responses to predictors. The corpus here closely parallels that of Brodbeck et al. (2022), with the same protocols and stimuli as native speakers, allowing for detailed comparisons between native and second-language speakers. Furthermore, our dataset includes two second-language groups of typologically distinct native languages (Sinhala and Mandarin) listening to the same text, affording the investigation of native-language specific transfer effects on phonological, morphological, and syntactic processing. This corpus will provide vital data for further studies of the neural mechanisms involved processing second language speech.

## Acknowledgments

This work was supported by NIDCD NIH R01-DC019394 (JS), NSF SMA-1734892 (JS), NSF BSC-1749407 (EL) and a Seed Grant from the UMD Brain and Behavior Institute (EL, JS).

## Footnotes

a A note on terminology: recent work has suggested that classifying speakers into ‘native’ and ‘non-native’ groups obscures rich and systematic variation in language knowledge (Weissler et al., 2023). While we acknowledge this complexity, in this paper we operationalized the term ‘second-language learner’ as someone who did not start learning the second language before the age of 5, and who was not immersed in an environment of native speakers before the age of 12 — the criteria for our study.

## References

Bell, A. J., & Sejnowski, T. J. (1995). An Information-Maximization Approach to Blind Separation and Blind Deconvolution. Neural Computation, 7 (6), 1129–1159. doi: 10.1162/neco.1995.7.6.1129

Bhattasali, S., Fabre, M., Luh, W.-M., Al Saied, H., Constant, M., Pallier, C., . . . Hale, J. (2019). Localising memory retrieval and syntactic composition: an fMRI study of naturalistic language comprehension. *Language*, Cognition and Neuroscience, 34 (4), 491–510.

Brennan, J. R., & Hale, J. T. (2019). Hierarchical structure guides rapid linguistic predictions during naturalistic listening. PLOS ONE, 14 (1), e0207741. doi: 10.1371/journal.pone.0207741

Brodbeck, C. (2023). TRF-Tools. Retrieved from https://github.com/christianbrodbeck/TRF-Tools

Brodbeck, C., Bhattasali, S., Cruz Heredia, A. A., Resnik, P., Simon, J. Z., & Lau, E. (2022). Parallel processing in speech perception with local and global representations of linguistic context. eLife, 11, e72056. doi: 10.7554/eLife.72056

Brysbaert, M., & New, B. (2009). Moving beyond Kučera and Francis: A critical evaluation of current word frequency norms and the introduction of a new and improved word frequency measure for American English. Behavior Research Methods, 41 (4), 977–990. doi: 10.3758/BRM.41.4.977

Clahsen, H., & Felser, C. (2006a). Continuity and shallow structures in language processing. Applied Psycholinguistics, 27 (1), 107–126. doi: 10.1017/S0142716406060206

Clahsen, H., & Felser, C. (2006b). Grammatical processing in language learners. Applied Psycholinguistics, 27 (1), 3–42. doi: 10.1017/S0142716406060024

Clahsen, H., & Felser, C. (2006c). How native-like is non-native language processing? Trends in Cognitive Sciences, 10 (12), 564–570. doi: 10.1016/j.tics.2006.10.002

Crosse, M. J., Di Liberto, G. M., Bednar, A., & Lalor, E. C. (2016). The multivariate temporal response function (mtrf) toolbox: A matlab toolbox for relating neural signals to continuous stimuli. Frontiers in Human Neuroscience, 10 . doi: 10.3389/fnhum.2016.00604

Davies, M. (2017). Corpus of Contemporary American English (COCA). Harvard Dataverse. doi: 10.7910/DVN/AMUDUW

Desikan, R. S., Ségonne, F., Fischl, B., Quinn, B. T., Dickerson, B. C., Blacker, D., . . . Killiany, R. J. (2006). An automated labeling system for subdividing the human cerebral cortex on MRI scans into gyral based regions of interest. NeuroImage, 31 (3), 968–980. doi: 10.1016/j.neuroimage.2006.01.021

Di Liberto, G. M., Nie, J., Yeaton, J., Khalighinejad, B., Shamma, S. A., & Mesgarani, N. (2021). Neural representation of linguistic feature hierarchy reflects second-language proficiency. NeuroImage, 227, 117586. doi: 10.1016/j.neuroimage.2020.117586

Ferreira, F., & Chantavarin, S. (2018). Integration and Prediction in Language Processing: A Synthesis of Old and New. Current Directions in Psychological Science, 27 (6), 443–448. doi: 10.1177/0963721418794491

Fischl, B. (2012). FreeSurfer. NeuroImage, 62 (2), 774–781. doi: 10.1016/j.neuroimage.2012.01.021

Gramfort, A., Luessi, M., Larson, E., Engemann, D. A., Strohmeier, D., Brodbeck, C., . . . Hämäläinen, M. S. (2014). MNE software for processing MEG and EEG data. NeuroImage, 86, 446–460. doi: 10.1016/j.neuroimage.2013.10.027

Hahne, A. (2001). What’s Different in Second-Language Processing? Evidence from Event-Related Brain Potentials. Journal of Psycholinguistic Research, 30 (3), 251–266. doi: 10.1023/A:1010490917575

Hahne, A., & Friederici, A. D. (2001). Processing a second language: late learners’ comprehension mechanisms as revealed by event-related brain potentials. Bilingualism: Language and Cognition, 4 (2), 123–141. doi: 10.1017/S1366728901000232

Heafield, K. (2011). KenLM: Faster and Smaller Language Model Queries. In Proceedings of the Sixth Workshop on Statistical Machine Translation (pp. 187–197). Edinburgh, Scotland: Association for Computational Linguistics.

Kaan, E. (2014). Predictive sentence processing in L2 and L1: What is different?* :. Linguistic Approaches to Bilingualism, 4 (2), 257–282. doi: 10.1075/lab.4.2.05kaa

Lew-Williams, C., & Fernald, A. (2010). Real-time processing of gender-marked articles by native and non-native Spanish speakers. Journal of memory and language, 63 (4), 447–464. doi: 10.1016/j.jml.2010.07.003

McAuliffe, M., Socolof, M., Mihuc, S., Wagner, M., & Sonderegger, M. (2017). Montreal Forced Aligner: Trainable Text-Speech Alignment Using Kaldi. In Interspeech 2017 (pp. 498–502). ISCA. doi: 10.21437/Interspeech.2017-1386

McLaughlin, J., Osterhout, L., & Kim, A. (2004). Neural correlates of second-language word learning: minimal instruction produces rapid change. Nature Neuroscience, 7 (7), 703–704. doi: 10.1038/nn1264

McLaughlin, J., Tanner, D., Pitkänen, I., Frenck-Mestre, C., Inoue, K., Valentine, G., & Osterhout, L. (2010). Brain Potentials Reveal Discrete Stages of L2 Grammatical Learning. Language Learning, 60 (s2), 123–150. doi: 10.1111/j.1467-9922.2010.00604.x

Oldfield, R. C. (1971). The assessment and analysis of handedness: the Edinburgh inventory. Neuropsychologia, 9 (1), 97–113. doi: 10.1016/0028-3932(71)90067-4

Pollan, M. (2002). The Botany of Desire: A Plant’s-Eye View of the World. Random House Publishing Group.

Smith, S. M., & Nichols, T. E. (2009). Threshold-free cluster enhancement: Addressing problems of smoothing, threshold dependence and localisation in cluster inference. NeuroImage, 44 (1), 83–98. doi: 10.1016/j.neuroimage.2008.03.061

Taulu, S., & Simola, J. (2006). Spatiotemporal signal space separation method for rejecting nearby interference in MEG measurements. Physics in Medicine & Biology, 51 (7), 1759. doi: 10.1088/0031-9155/51/7/008

Weissler, R. E., Drake, S., Kampf, K., Diantoro, C., Foster, K., Kirkpatrick, A., . . . Baese-Berk, M. M. (2023). Examining linguistic and experimenter biases through “non-native” versus “native” speech. Applied Psycholinguistics, 1–15. doi: 10.1017/S0142716423000115

